# Homo-Oligomerisation in Signal Transduction: Dynamics, Homeostasis, Ultrasensitivity, Bistability

**DOI:** 10.1101/758789

**Authors:** Daniel Koch

**Affiliations:** Randall Centre for Cell & Molecular Biophysics, King’s College London, London SE1 1UL, United Kingdom

## Abstract

Homo-oligomerisation of proteins is a ubiquitous phenomenon whose exact role remains unclear in many cases. To identify novel functions, this paper provides an exploration of general dynamical mathematical models of homo-oligomerisation. Simulation and analysis of these models show that homo-oligomerisation on its own allows for a remarkable variety of complex dynamic and steady-state regulatory behaviour such as transient overshoots or homeostatic control of monomer concentration. If post-translational modifications are considered, however, conventional mass-action kinetics leads to thermodynamic inconsistencies due to asymmetric combinatorial expansion of reaction routes. Introducing a conservation principle to balance rate equations re-establishes thermodynamic consistency. Using such balanced models it is shown that oligomerisation can lead to bistability by enabling pseudo-multisite modification and kinetic pseudo-cooperativity via multi-enzyme regulation, thereby constituting a novel motif for bistable modification reactions. Due to these potential signal processing capabilities, homo-oligomerisation could play far more versatile roles in signal transduction than previously appreciated.

## Introduction

Homo-oligomerisation of proteins, i.e. the assembly of supramolecular protein complexes made up from multiple identical subunits, is a ubiquitous phenomenon. In vertebrates, about 30-50% of all proteins form homo-oligomers consisting of two or more subunits [1, 2]. Oligomerisation may offer several advantages: it can be a way to economically assemble larger structures (thereby reducing genome size) and allows for a higher error-free transcription chance for individual subunits. Moreover, it can provide additional regulatory control via allostery and cooperative binding events (hemoglobin being the classical example) [3, 4]. Yet, in many cases the function of homooligomerisation remains unclear.

Dynamical mathematical models based on ordinary differential equations (ODEs) have been extensively used to study many important motifs, mechanisms and phenomena in signal transduction networks. To lesser extent, ODE models have also been used to study signal transduction processes involving homo-oligomers. Such theoretical studies have shown that in addition to the well-known role in the emergence of ultrasensitive responses via cooperative binding, oligomerisation can provide an additional layer of control over such responses. Bouhaddou an Birtwistle, for instance, showed that different oligomerisation routes provide an effective means of tuning ultrasensitive, cooperative responses [5]. Buchler and Louis showed that homo-oligomerisation itself can lead to modest ultra-sensitivity independently from cooperativity [6]. If coupled to positive feedback, the ultrasensitivity generated e.g. by homo-dimerisation is sufficient for the emergence of bistability [7]. For signalling involving dimeric receptors and substrate activation, the presence of a single/dual activation mechanism can lead to complex, non-linear signal dynamics [8]. Taken together, this highlights the importance of homo-oligomerisation and the use of mathematical modelling as a tool to study their roles in signal transduction.

However, above mentioned studies focussed on specific questions, contexts or systems. A general analysis of homo-oligomerisation in terms of assembly dynamics, steady state behaviour and the potential effects of post-translational modifications (PTMs) is neither covered by classical and popular textbooks on mathematical or system’s biology (see e.g. 9-[13]), nor is the author aware of such an analysis in the recent research literature. It thus seems that an exploration of general dynamical mathematical models of homooligomerisation and is still lacking. This paper provides such an exploration. As this study focusses solely on homo-oligomerisation, we will leave out the prefix ‘homo-’ in the remainder of this article for the sake of brevity.

Starting with simple mass action kinetics based models of dimerisation to tetramerisation, we will study assembly dynamics and steady state behaviour numerically. Although the first presented models are very simple, it is found that they are capable of complex dynamical and steady state behaviour such as undulations and homeostatic regulation.

Next, PTMs of oligomers are considered. Surprisingly, the application of conventional mass action rate laws easily results in thermodynamically inconsistent models due to combinatorial expansion of the oligomerisation routes upon modification. The issue can be circumvented by balancing the rate expressions based on a rate conservation principle.

Finally, two novel mechanisms based on oligomerisation leading to ultrasensitive, bistable PTM responses will be presented: pseudo-multisite modification and regulation by multiple enzymes.

The focus of the current work is to demonstrate that oligomerisation enables more complex regulatory behaviour than previously appreciated. While the broad scope of a general analysis of dynamical mathematical models of oligomerisation does not permit an exhaustive treatment of all aspects within the limit of a single article, some of the most important implications and avenues for future research will be outlined in the discussion.

## Results

### Simple mass action models of oligomerisation: ultrasensitivity and homeostasis

Let us begin by assuming that a general protein *A* can form symmetric oligomers with a maximum number of *n* subunits (protomers) per oligomeric complex. We furthermore assume that *A* can form all intermediate oligomeric species with *m* subunits (where *m ϵ* ℕ, 1 < *m* < *n*) and that each oligomeric species is formed through simple one-step, second-order binding reactions described by mass action kinetics. For the remainder of this article, we will study oligomers with a maximum of four protomers or less, i.e. tetramers, trimers and dimers. It is likely that many of the presented findings apply to higher-order oligomers as well.

In the case of tetramers we therefore assume that tetramers can be formed by the association of two dimers or, alternatively, of a trimer and a monomer. The reaction scheme and the reaction rates *v*_*i*_ for the individual reactions can then be summarised as in Figure 1. Denoting the monomeric to tetrameric species by *A*, …, *AAAA* we can now formulate the system’s equations:

**Figure 1.**
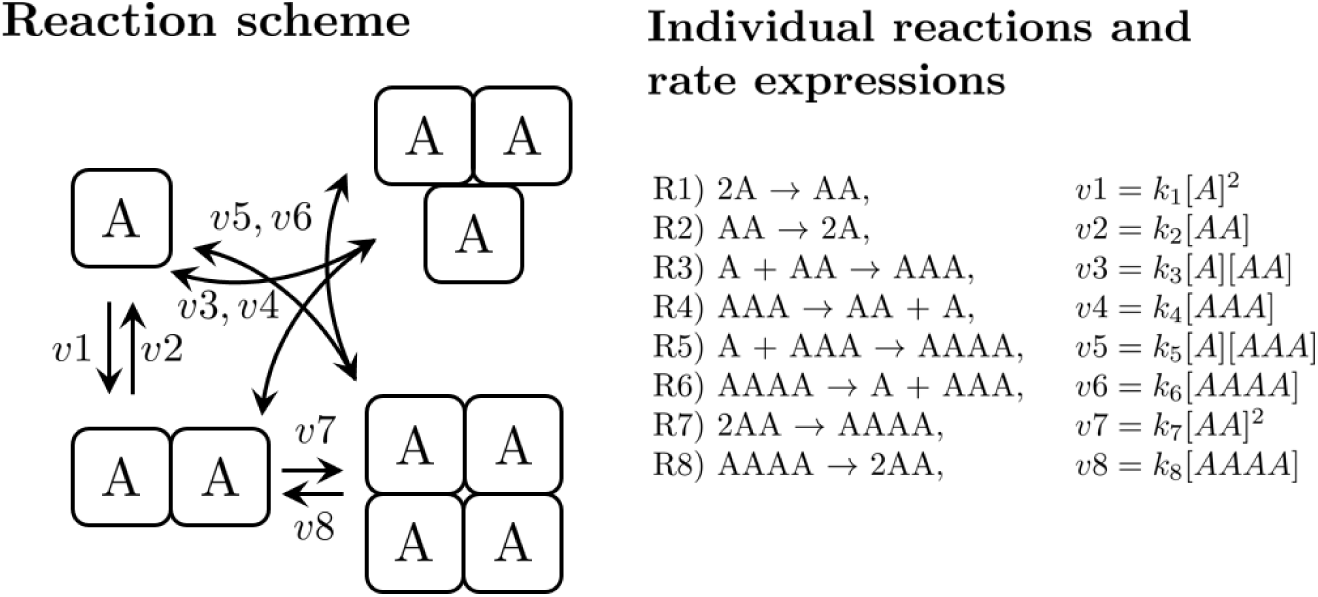
Model scheme of homo-tetramerisation based on conventional mass action kinetics assuming that all intermediate species (dimers and trimers) are possible in the reaction pathway. See text for the differential equations describing the system.

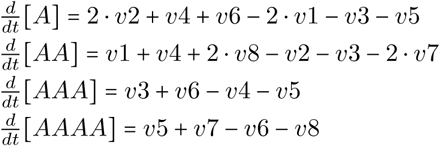

The total amount of subunits is conserved by the relation *A*_*tot*_ = [*A*] + 2 · [*AA*] + 3 · [*AAA*] + 4 · [*AAAA*] and can be used to eliminate one of the above equations. Note that models for tri- or dimerisation can be obtained simply by removing reactions R5-R8 or R3-R8, respectively. For the sake of simplicity, we will begin by assuming equal rate constants of 10^6^*mol* ^−1^*s*^−1^ for all association reactions and equal rate constants of 0.1*s*^−1^ for all dissociation reactions, thereby yielding a *K*_*d*_ value of 0.1 *µM* for all reactions, a typical value for many protein-protein interactions.

Time course simulations of the system with initial conditions [*A*]_0_ = *A*_*tot*_ = 10*µM* show association dynamics typical for binary protein-protein interactions in the case of dimerisation, whereas trimerisation and tetramerisation reactions exhibit a transient overshoot of dimers followed by a slower decrease and increase in trimers and tetramers, respectively (Figure 2A-C). The amplitude and position of such overshoots depend strongly on the monomer concentration at the beginning of the reaction (Figure 2C, inset). More complex dynamics such as dampened oscillations/undulations on different timescales are possible (Figure 2B, inset). If the individual oligomeric species possess different biological functionality, such dynamics could be a mechanism for dynamic signal encoding as will be outlined in the discussion in more detail.

**Figure 2.**
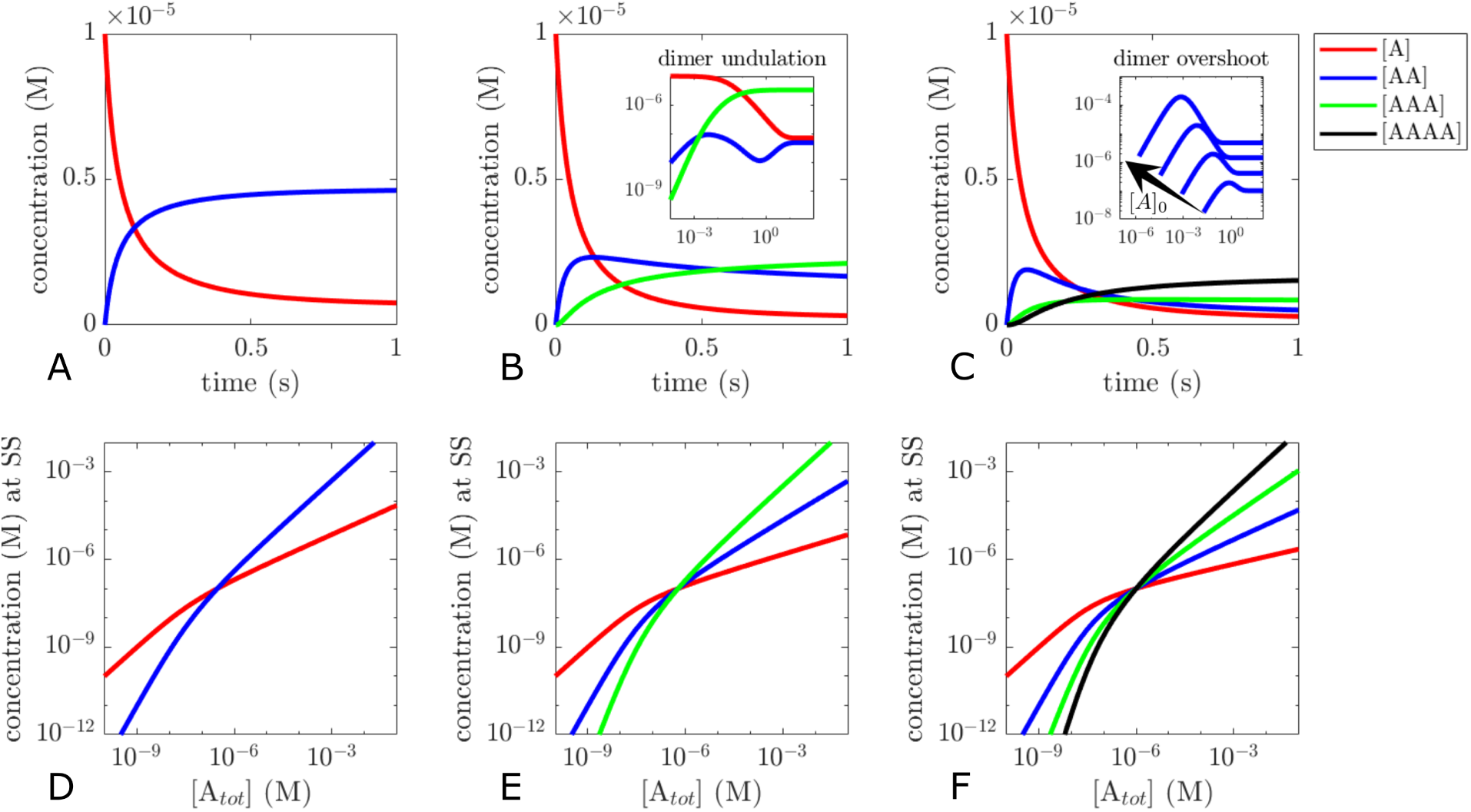
Time course simulations (A-C) and steady state analysis (D-F) for dimers (A,D), trimers (B,E) and tetramers (C,F). Initial conditions for A-C [*A*]_0_ = 10*µM*, [*AA*]_0_ = [*AAA*]_0_ = [*AAAA*]_0_ = 0*M*. Parameters (A and D) *k*_1_ = 10^6^*mol*^−1^ *s*^−1^, *k*_2_ = 0.1*s*^−1^, (B and E) *k*_1_ = *k*_3_ = 10^6^*mol*^−1^*s*^−1^, *k*_2_ = *k*_4_ = 0.1*s*^−1^, (C and F) *k*_1_ = *k*_3_ = *k*_5_ = *k*_7_ = 10^6^*mol*^−1^*s*^−1^ (C inset *k*_3_ = 10^8^*mol*^−1^*s*^−1^), *k*_2_ = *k*_4_ = *k*_6_ = *k*_8_ = 0.1*s*^−1^.

Numerical steady state analysis shows that the dose-response curves for the individual species meet in a single intersection point (Figure 2D-F), mirroring the assumption that all reactions have the same *K*_*d*_ value. Local sensitivity analysis at *A*_*tot*_ = 10*nM* with 2% perturbation yields relative sensitivities 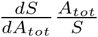(where *S* is the steady state concentration of either *A*, …, *AAAA*) of 0.87 for monomers and 1.76 for dimers in the dimerisation model, 2.58 for trimers in the trimersation model and 3.46 for tetramers in the tetramerisation model. The analysis confirms again that oligomerisation at concentrations below the *K*_*d*_ can lead to modest ultrasensitivity in response to changes in total protein concentration, and that ultrasensitivity can increase with higher number of protomers per complex (as can also be seen from the increasing slopes in Figure 2E,F) [6].

For many proteins able to form higher order oligomers, the presence of a single or a small subfraction of possible oligomeric species often dominates over other potential intermediate species [14], indicating that oligomerisation is often cooperative and that *K*_*d*_ values differ for the individual reactions. Varying the model parameters in a way to favour formation of the highest order oligomer in the trimerisation and tetramerisation model (e.g. by increasing association rate constants) can reproduce the dominance of the highest order oligomer over large concentration ranges (Figure 3A,B). This also leads to a shift of intersection points, resulting in different apparent *K*_*d*_ values between the individual intermediate oligomerisation reactions. Tweaking of the parameters allows to shift the curves for each individual species into almost any direction (data not shown). sParameter variation also highlights the flipside of the coin of oligomeric ultra-sensitivity. If we consider the monomer concentration at higher total protein concentrations in the inset of Figure 3B, it becomes apparent that oligomerisation can be an efficient homeostatic regulatory mechanism of the monomer concentration (relative sensitivity of 0.25 for monomers at *A*_*tot*_ = 100*µM*). This would be plausible in situations where monomers are the biologically active species. Note that this mechanism does not require a complex feedback organisation typically associated with homeostasis [15, 16].

**Figure 3.**
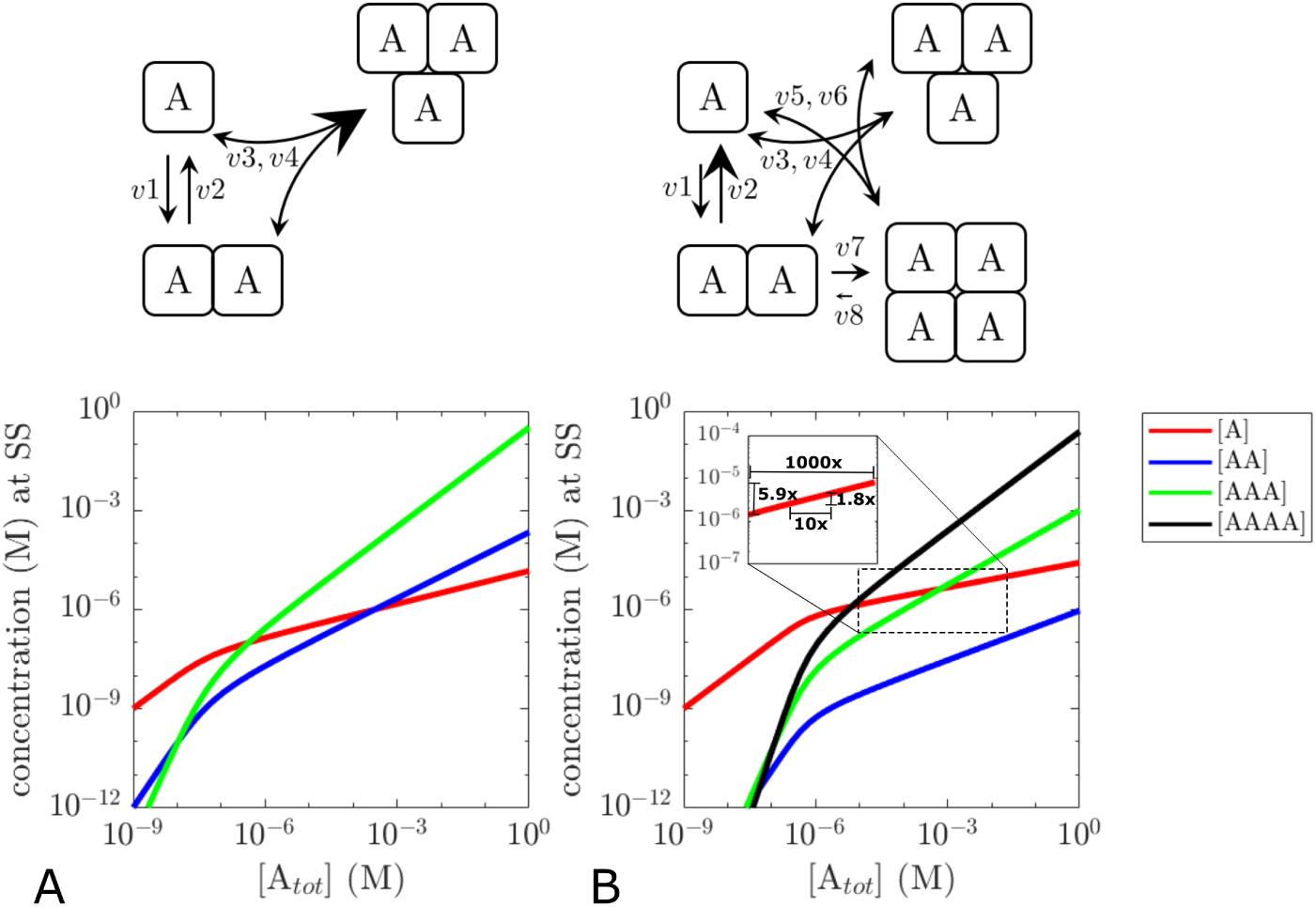
Steady state analysis of trimerisation and tetramerisation models with varied parameters. The relative change of parameters is visualised in the upper reaction schemes. Parameters (A) *k*_1_ = 10^6^ *mol*^−1^*s*^−1^, *k*_3_ = 10^8^*mol*^−1^ *s*^−1^, *k*_2_ = *k*_4_ = 0.1*s*^−1^, (B) *k*_1_ = 3.2 × 10^6^*mol*^−1^*s*^−1^, *k*_2_ = 2400*s*^−1^ *k*_3_ = 3.45 × 10^6^*mol*^−1^*s*^−1^, *k*_4_ = 0.083*s*^−1^, *k*_5_ = 4.8 × 10^6^*mol*^−1^*s*^−1^, *k*_6_ = 0.525*s*^−1^, *k*_7_ = 3 × 10^6^ *mol*^−1^ *s*^−1^, *k*_8_ = 1.0525 × 10^−5^*s*^−1^.

### Considering PTMs: ensuring thermodynamical consistency

Just like non-oligomeric proteins, oligomeric proteins are subject to various post-translational modifications such as phosphorylation, ubquitinylation, lipidation and others. Sometimes these modifications can regulate the equilibria between monomeric and oligomeric species via conformational changes or sterical hindrance. Other times these modifications are irrelevant to the protein’s oligomerisation behaviour. As we will see shortly, even accounting merely for a single PTM makes model formulation of anything higher than dimers unlikely more difficult due to the combinatorial expansion of potential oligomerisation routes. In the following, we will therefore restrict mass action models to a maximum of trimerisation.

Unfortunately, combinatorial expansion is not the only challenge when PTMs of oligomers are considered: it is remarkably easy to slip into thermodynamical inconsistency even with models based purely on conventional mass action kinetics. Before we incorporate PTMs into our models we will therefore formulate some biochemical intuitions and expectations which will later guide us to avoid thermodynamic inconsistency.

Let us suppose an oligomeric protein can be modified by a PTM at a single site. For the sake of simplicity we assume that the site lies remote from the oligomerisation interface and does not alter any of the reaction parameters. Intuitively, we would then expect the following to be true:

(a) The association rate constants (a1), dissociation rate constants (a2) and therefore equilibrium and dissociation constants (a3) are identical for the following oligomerisation reactions:
  - only unmodified protein
  - only modified protein
  - modified with unmodified protein
(b) If the modifying enzyme is added to an equilibrated mixture of completely unmodified monomers and oligomers, both monomers and oligomers will be modified over time, yet the total concentrations of monomers and oligomers remains constant.
(c) There are 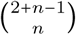 possibilities to combine unmodified/modified monomers into symmetric n-tamers. Thus, if equal parts of unmodified/modified monxomers are mixed to form n-tamers, the isoform distribution of n-tamers with 0 ≤ *i* ≤ *n* modified subunits will be binomal.

To proceed with model formulation, let *A*\*, *AA*\*, *AA*\*\*, … denote modified monomers, dimers with one and dimers with two modified protomers and so forth. Due to assumed symmetry, molecules such as *A*\* *A* and *AA*\* are identical. Keeping the assumption that each oligomeric species is formed through binary association reactions, it follows from (a1)-(c1) that each pair of molecular species X,Y which is able to associate in the absence of any PTMs is also able to associate with identical reaction parameters regardless of how many protomers of X or Y are modified. We will assume that there is a modifying enzyme *E*1 and a demodifying enzyme *E*2 which operate by a non-cooperative, irreversible and distributive mechanism and that all molecular species, regardless the number of their protomers, are (de-)modified with the same kinetic parameters, i.e. the oligomeric state does not influence the (de-)modification reactions. These assumptions reflect the situation where a PTM does not induce conformational changes and lies remote from the oligomerisation interface, allowing the enzymes to access the PTM site equally in all oligomeric species. We therefore expect the individual monomeric and oligomeric species to compete for enzymes *E*1 and *E*2. In situations with multiple competing substrates *S*_1_, …, *S*_*n*_ an irreversible Michaelis-Menten type rate law of the form:

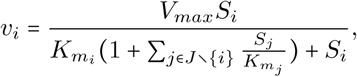

where *J* = {1, …, *n*}, can be employed to describe the rate of consumption *v*_*i*_ of substrate *S*_*i*_ [17]. That is, the individual substrates act as competitive inhibitors for each other. We are now able to formulate reaction schemes, reaction rates and model equations.

Figure 4A shows the reaction scheme and rate expressions for the dimerisation model based on mass action kinetics for oligomerisation and mentioned Michaelis-Menten type rate law for addition and removal of PTMs. The equations are:

**Figure 4.**
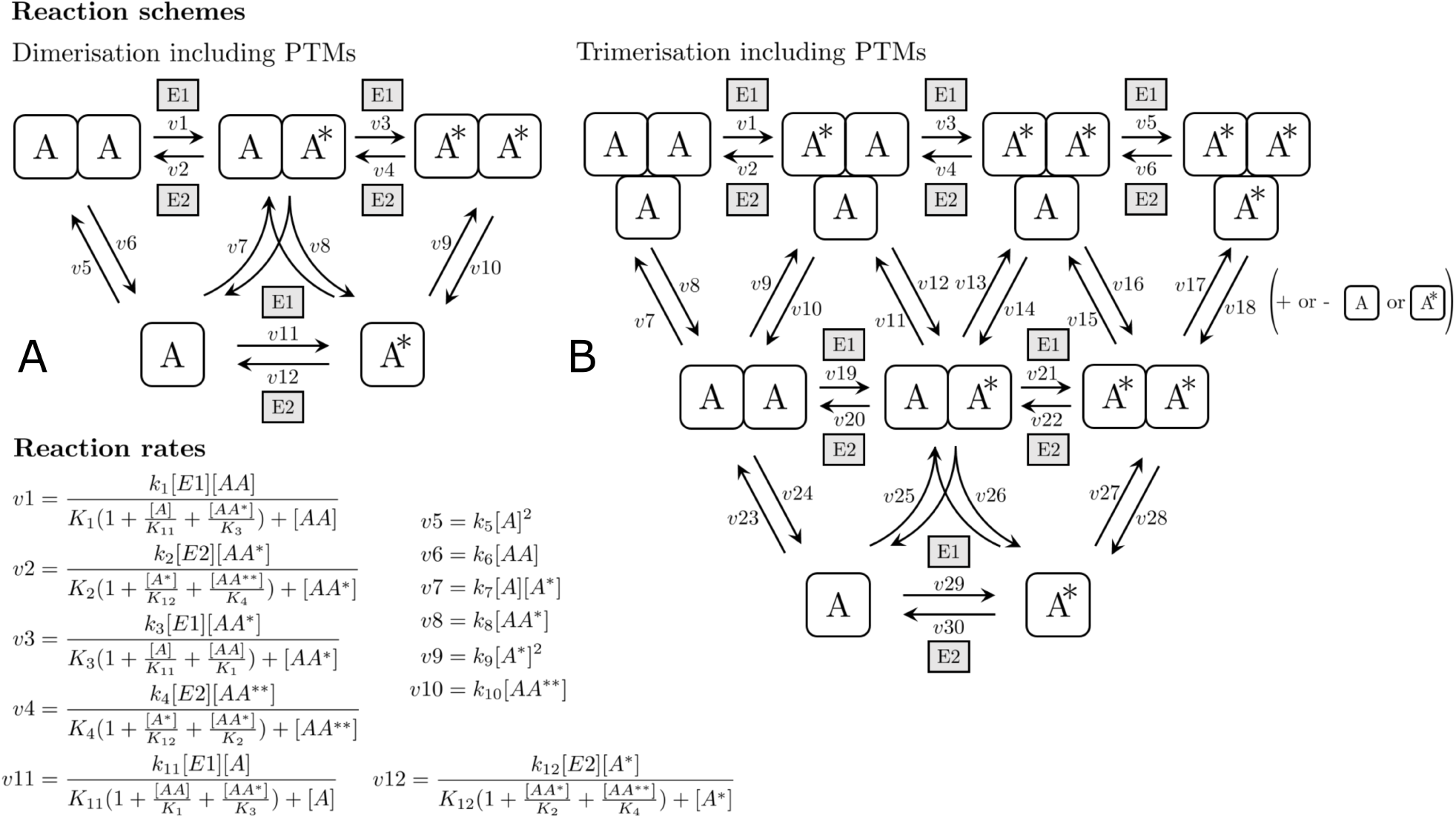
Reaction schemes for the mass action kinetics models of dimerisation (A) and trimerisation (B) including reversible post-translational modifications. See supplementary information for reaction rates and ODEs of the trimersation model.

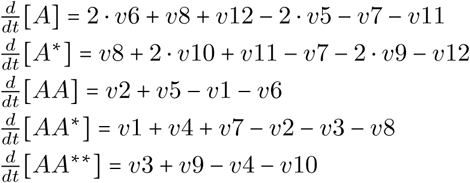

Figure 4B shows the reaction scheme for the trimerisation model including PTMs. See supplementary section 1 for reaction rates and model equations. Note that the reaction scheme for trimerisation and a single PTM is already more complex than the tetramerisation scheme without PTMs. For dimerisation, we will use the same parameter values as in Figure 2A. For trimerisation, we will use the same parameter values as in Figure 3A. We chose catalytic rate constants to be *k*_*cat*_ *=* 1*s*^−1^ and Michaelis constants to be *K*_*m*_ *=* 1*µM* for all (de-)modification reactions in both models.

Let us first consider the effects of adding catalytic concentrations of E1 to an equilibrated mixture of monomers and oligomers. Interestingly, as the modification reaction proceeds, transient changes in the total concentrations of monomeric and oligomeric species occur (Figure 5A; the total concentrations being the sum of concentrations of all modified isoforms for a given monomeric or oligomeric species). The amplitude and direction of these transients appear to be both influenced by the maximum number of protomers. Intuitively, the phenomenon results from the accessibility of additional oligomerisation routes during the modification reaction. Consider for example the dimerisation reaction scheme (Figure 4A). If *A* is either completely unmodified or modified, a single reaction route between a monomeric and a dimeric species is available. If a mixture of *A* and *A*\* is present, a third route appears: the reaction between *A* and *A*\* to *AA*\*. We would therefore expect that decreasing the velocity of oligomerisation/dissociation or increasing the speed of modification would diminish the effect, as both changes would limit *A*’s capacity to change its oligomerisation state during the transition time from completely unmodified protein to complete modification. Indeed, while decreasing the velocity of oligomerisation/dissociation reduces the amplitude, increasing the enzyme concentration limits the peak width of the amplitude (supplementary Figure S1).

**Figure 5.**
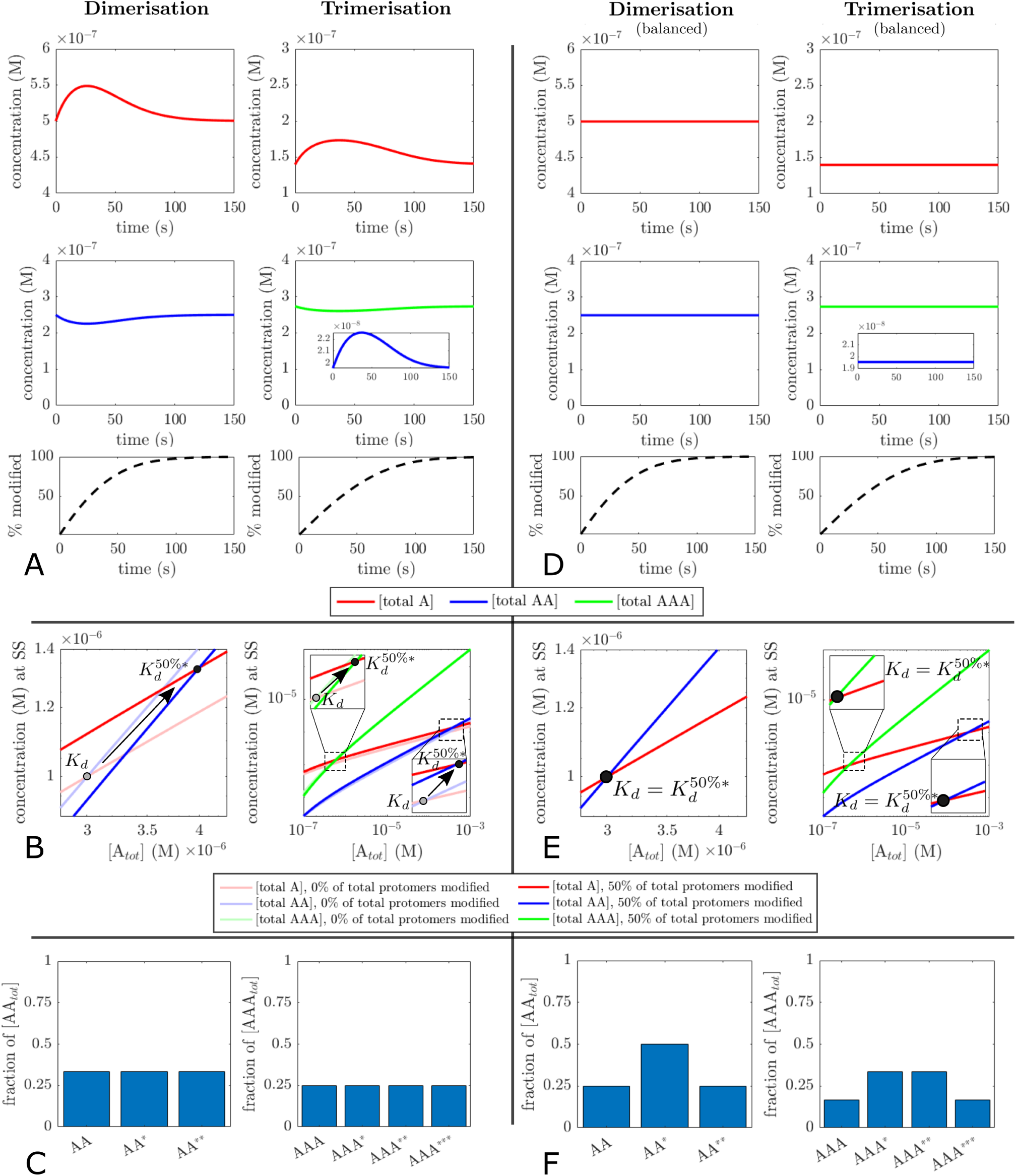
Behaviour of di- and trimerisation models upon post-translational modification. (A,D) timecourses of total monomer and total oligomer concentrations after addition of 50*nM* [*E*1] starting with [*A*_*tot*_] = 10*µM* at equilibrium. Dimerisation rate constants: 10^7^*mol* ^−1^*s* ^−1^, trimerisation rate constants: 10^6^*mol*^−1^*s*^−1^, dissociation rate constants of dimers and trimers 10*s*^−1^. Black-dotted curves show progress of the modification reaction. (B,E) apparent dissociation constants and (C,F) distribution of PTM isoforms of the highest order oligomer in equimolar mixtures of modified and unmodified protein at equilibrium.

Clearly, these transient changes contradict expectation (b) and no experimental data so far (as far as the author is aware) reported transient alterations of the oligomerisation state upon post-translation without affecting association/dissociation rates. To further explore the models and to test whether the transient changes could be a modelling artefact, let us see what happens if equimolar mixtures of modified and unmodified *A* are simulated to steady state. According to (a3) we would expect *K*_*d*_ values to be identical to that of completely unmodified and completely modified *A*. We would furthermore expect a binomial distribution of modified oligomers according to (c). However, both models show altered *K*_*d*_ values (Figure 5B) and a uniform distribution of modified oligomers (Figure 5C). Taken together, this suggests that the transient changes are indeed a modelling artefact resulting from thermodynamic inconsistency.

Where did we go astray if using simple mass action kinetics, arguably the most established deterministic modelling approach for chemical reactions, leads to thermodynamic inconsistency? The answer is: naively deriving ordinary differential equations by applying mass action kinetics and stoichiometrical balancing to reaction schemes such as in Figure 4. To circumvent this issue, we need to introduce a more general notation for oligomeric species and formulate a further expectation.

Let 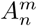 denote a *m*-times modified n-tamer, i.e. an oligomeric complex with *n* ϵ ℕ^≥1^ identical subunits of which 0 ≤ *m* ≤ *n* carry a PTM at site *x*. Let *j* ϵ ℕ^≥1^ be the number of reversible bimolecular association reactions between lower-order complexes 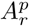 and 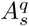 able to form 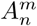, where *r* + *s* = *n*.

Next, let *I* (*A*_*n*_) = {{*r*_1_, *s*_1_}, {*r*_2_, *s*_2_}, …} define the set of actually occurring combinations of oligomeric orders of *A*_*n*_’s educts 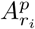 and 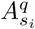, where *p* + *q* ≤ *n* and *r*_*i*_ + *s*_*i*_ *= n*. For example, if a hexamer can be assembled from dimers with tetramers and from monomers with pentamers (of any PTM status), *I*(*A*_6_) would be the set {{2, 4}, {1, 5}}.

From (a1) and (a2) it follows that all forward steps between 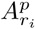 and 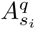 producing 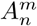 have the same association rate constant *k*_*i*_ and each corresponding reverse step has the same dissociation rate constant 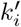 for all possible values of *m, p* and *q*. Unaltered rate constants imply unaltered equilibrium constants. Therefore, we expect that

(d) the PTM status is irrelevant for the equilibrium. Thus, the equilibrium is solely determined by the total concentrations of each n-tamer, i.e. the sum of all PTM-isoforms of *A*_*n*_:

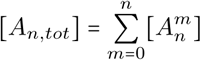

In other words: from the perspective of the oligomerisation equilibrium, all PTM isoforms of a n-tamer are treated as a single species.

For a graphical illustration of the definitions and the equilibrium situation see supplementary Figure S2. We can now formulate the following principle:

### Conservation of oligomerisation rates

If a PTM does not influence the parameters of an oligomerisation reaction, the sum of all rates *v*_*i*_, 1 ≤ *i* ≤ *j*, of reactions leading to a n-tamer *A*_*n*_ of any modification status is equal to the association rates based on the total concentrations of *A*_*n*_’s educts (i.e. all modification isoforms). Conversely, the sum of all rates 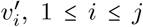, of reactions dissociating *A*_*n*_ of any modification status is equal to the dissociation rate based on *A*_*n*_’s total concentration. That is, at all times

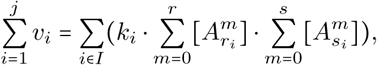

and

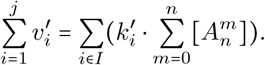

We call the right-hand side of each identitiy the *efective rates*.

The principle’s name is chosen in analogy to the conservation of mass as it conserves the reaction rates at given total concentrations of educts regardless the distribution of PTM isoforms. Although it might appear complicated, it is straightforward to illustrate the principle using dimerisation as an example. Applying it to Figure 4A yields *I* (*A*_2_) = {{1, 1}} and thus

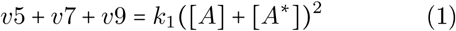

and

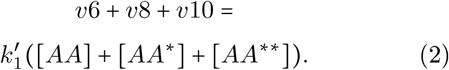

How does this principle help us to avoid thermodynamic inconsistency? The principle relates the *j* reaction rates from the reaction scheme (left-hand side (LHS) of (1) and (2)) to the thermodynamically expected effective rates (right-hand side (RHS) of (1) and (2)). Instead of assigning a *priori* rate expressions based on mass action kinetics, we will assign rates only after making sure the principle is not violated. In the first step of this check we will expand the RHS of the rate conservation identity. In a second step, we will substitute the rates on the LHS with their mass action kinetics expression. Next, we will compare both sites: if they are equal, all rates can readily be identified with their mass action kinetics expression. If they are not equal, we balance the terms on the LHS where the discrepancy occurs by introducing coefficients that ensure the validity of the identity. Lastly, we identify the respective rates with the balanced terms. Let us illustrate this using equations (1) and (2) derived from the dimerisation scheme:

(1) → *expanding RHS, substituting LHS*:

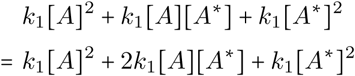
→ *balancing deviating terms in LHS:*

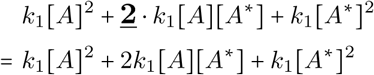
→ *assigning reaction rates:*

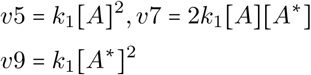
(2) → *expanding RHS, substituting LHS:*

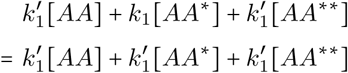
→ *no balancing necessary*
→ *assigning reaction rates:*

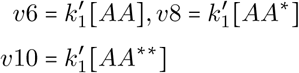

It is straightforward to apply the same procedure to the trimerisation model (cf. supplementary section 2). Other things being equal, the dimerisation and trimerisation model updated with the balanced rate expressions exhibit neither transient changes in the oligomerisation upon modification (Figure 5D), nor shifts in the apparent *K*_*d*_ of equimolar mixtures of unmodified/modified *A* (Figure 5E). The distribution of PTM isoforms at equilibrium is binomial (Figure 5F). Taken together, this indicates that the balanced models are indeed thermodynamically consistent.

The postulated rate conservation principle is no fundamental law and can readily be proven using the principle of detailed balance [18] as lemma (cf. supplementary section 3).

For a more intuitive understanding of this balancing procedure, it might be helpful to appreciate its similarity to stoichiometric balancing. Consider, for example, the chemical equation for the reaction of oxygen with hydrogen to water:

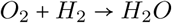

As the reader will have spotted, there are two oxygen atoms on the LHS, but only one on the RHS of the equation. As this violates the law of conservation of mass, we need to balance the equation by adding stoichiometric coefficients:

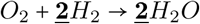

The rate balancing procedure presented in this paper is conceptually identical: As the PTM is assumed not to influence oligomerisation, we know that total formation and dissociation rates of oligomers are conserved; they must be the same as for unmodified protein. However, the PTM increases the number of possible oligomeric species due to combinatorial expansion. This expansion is asymmetric because there are more possibilities to combine modified and unmodified subunits to a n-tamer the higher its order: two for a monomer, three for a dimer, four for a trimer and so forth (cf. Figure 4B). This creates additional oligomerisation routes and alters the net-rates of oligomer formation and dissociation, thereby violating the rate conservation principle. To avoid this, we need to balance this purely combinatorial effect by introducing balancing coefficients for the rate expressions.

### Considering PTMs: ultrasensitivity and bistability

Now that we have thermodynamically consistent models of oligomers which can be post-translationally (de-)modified, we will explore the steady state behaviour in the presence of (de-)modification enzymes *E*1, *E*2 using the dimer model as an example. The relative fraction of modified dimer and monomer shows pronounced ultrasensitivity in response to increasing concentrations of modifying enzyme *E*1 (Figure 6A).

**Figure 6.**
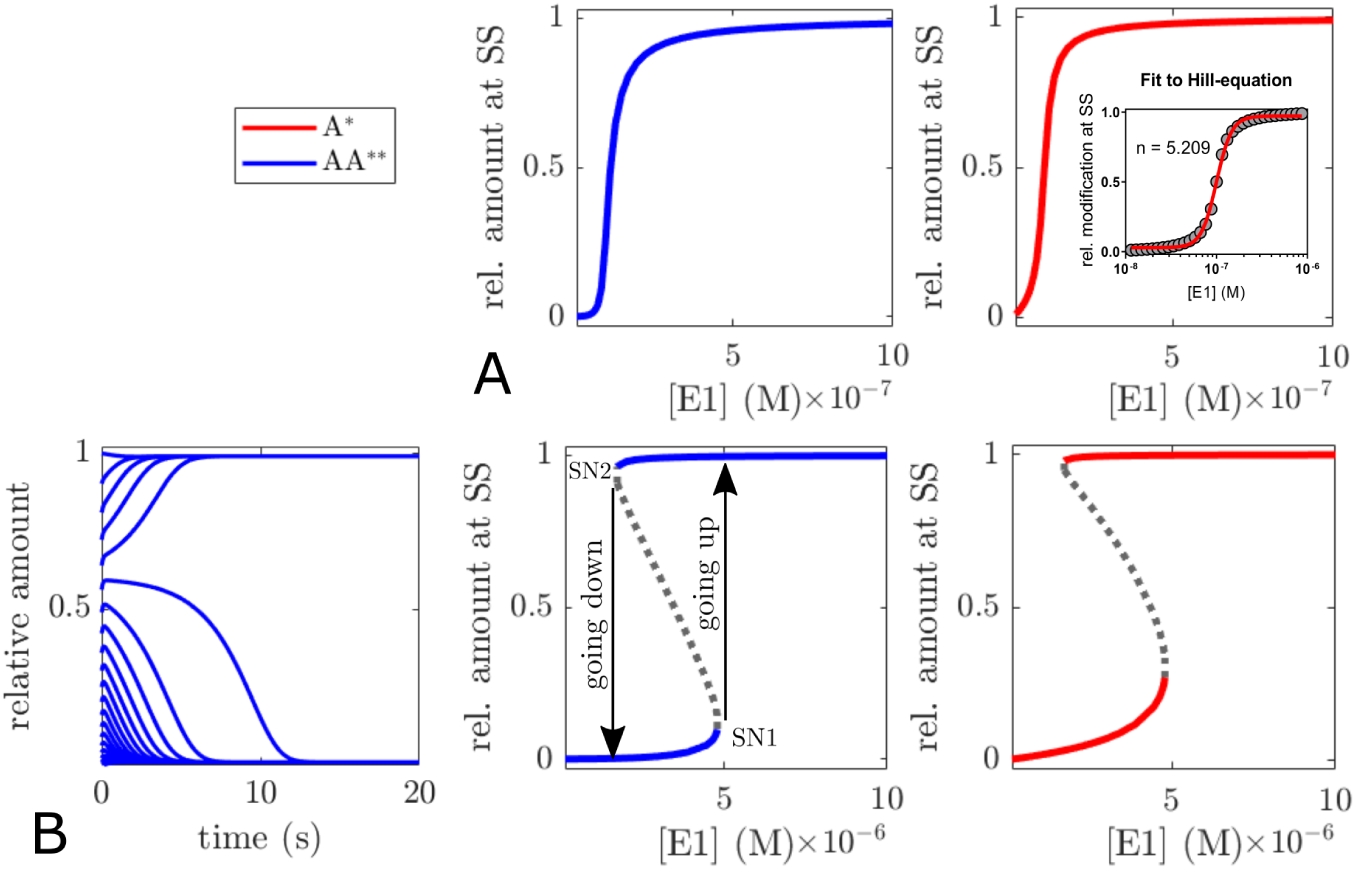
Ultrasensitivity and Bistability of modification response. (A) fractional modification of both monomers and dimers in response to increasing concentrations of modifying enzyme *E*1 is notably ultrasensitive. (B) left, time course simulations demonstrate that the approached steady state is determined by the initial conditions if demodification is assumed to be cooperative. Parameters *k*_2_ = 100*s* ^−1^, *k*_4_ = 10*s*^−1^, [*E*1] = 3*µM*, [*E*2] = 0.1*µM, A*_*tot*_ *=* 10*µM* with different fractional modification at t = 0), oligomerisation parameters as specified for Figure 5. (B) middle and right, bifurcation diagrams show identical parameter values for saddle node bifurcations of dimer and monomer modification. Unstable steady states are indicated by dotted lines.

On closer examination, this is not very surprising. Apart from some degree of zero-order ultrasensitivity arising from enzyme saturation [19], oligomerisation additionally creates a substrate competition situation between monomeric and oligomeric species for (de-)modification and provides pseudo-multisites for PTMs (i.e. multiple protomers with identical PTM sites). Both motifs are capable of generating ultra-sensitivity [20, 21]. Moreover, multisite modification can in principle generate bi- or multistability if there is a sufficient asymmetry in the sequential modification cycles, i.e. if either the demodification and/or the modification steps exhibit kinetic cooperativity [22-24]. For dual-site modification of monomeric proteins, Conradi and Mincheva have proven that in general, bistability must occur for some concentrations of demodifying and modifying enzyme if the product of the rate constants for the first modification and demodification steps is smaller than the product of the rate constants for the second modification and demodification steps [25]. Without considering the oligomeric nature, introducing positive kinetic cooperativity for the demodification of the dimer, i.e. assuming *k*_2_ > *k*_4_, would fulfill this requirement. Indeed, implementing this assumption leads to bistability with respect to the modification status in the dimer model (Figure 6B). As the bistable range increases with the number of cooperative modification steps [23], the likelihood for a bistable PTM status will increase with higher order oligomers.

Interestingly, not only the dimer, but also the monomer exhibits bistability even without multiple sites for PTMs. This becomes less surprising if one considers that the dimer is in equilibrium with the monomer allowing modified dimers to dissociate into monomers. Furthermore, when dimers are completely (de-)modified, substrate competition for (de-) modification of the monomer abates, allowing for more monomer (de-)modification.

While perhaps not uncommon, kinetic cooperativity might not be the only way to realise bistability in (pseudo-)multisite PTM systems. From a biochemical point of view, asymmetry in the (de-)modification rate of a multisite PTM protein could effectively be realised, too, if one of the (de-)modification steps would also be catalysed by another enzyme *E*3. Let us, for instance, assume that in a dually modified dimer, each PTM mutually prevents (e.g. due to sterical reasons) access to the other PTM for demodifying enzyme *E*3. Only when one of the PTMs has already been removed by demodifying enzyme *E*2 (which we assume to catalyse PTM removal from the singly and dually modified dimer equally well), can *E*3 bind to the singly modified dimer and catalyse the last demodification step. Assuming that *E*3 can also catalyse demodification of the modified monomer, the scheme for updated dimer model is shown in Figure 7A. Using the updated dimer model, it is not difficult to find parameter values that lead to bistability (Figure 7B), showing that multi-enzyme regulation can be an effective alternative for realising the asymmetry required for bistability in multisite PTM systems.

**Figure 7.**
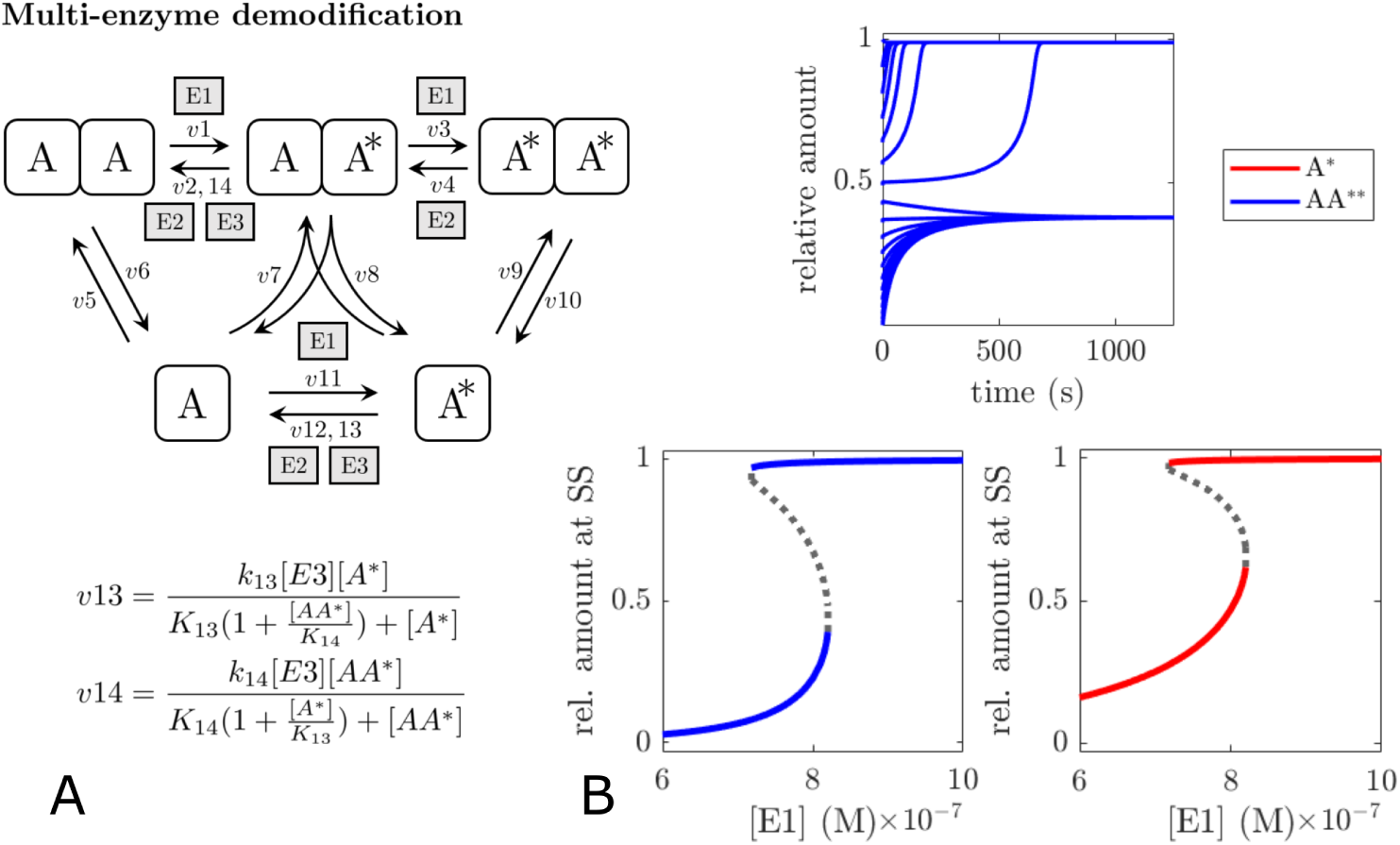
Bistability through multi-enzyme regulation of modification status of oligomers. (A) scheme of (balanced) dimer model with additional demodification enzyme *E*3 which can not catalyse the first step of the dimer demodification. (B) time course simulations and bifurcation plots demonstrating bistability in the dimer model. Parameters [*E*2] = 10*nM*, [*E*3] = 0.1*µM, k*_13_ = 0.1*s*^−1^, *k*_14_ = 100*s*^−1^, *K*_13_ = 10*µM, K*_14_= 1*µM*, oligomerisation parameters as specified for Figure 5.

## Discussion

### Complex dynamics and steady state behaviour

Even simple oligomerisation systems can in principle be capable of surprisingly complex behaviour. Dynamical phenomena such as transient overshoots of dimers followed by a slower increase in higher order oligomers will be relevant to proteins which are not in constitutive monomer/oligomer equilibrium. Examples include membrane receptors which oligomerise upon ligand binding [26] or proteins that oligomerise upon recruitment to a membrane. If dimers and higher order oligomers have different downstream signalling functions, such transients could be an effective way to encode the duration of the input signal (e.g. ligand presence or membrane recruitment) and thereby lead to different cellular responses for short and prolonged stimuli. The tumor suppressor p53 is a relevant example of a protein different biological activity for different oligomeric species [27]. As it is also involved in dynamic signal encoding leading to different cell-fate decisions [28], it is tempting to speculate that some of this could be the result of oligomerisation. Another promising candidate for dynamic signal encoding through oligomerisation could be the EGF-receptor for which dimers, trimers and tetramers have been described [29, 30]. A considerable list of higher-order homo-oligomers for which various intermediate forms have been observed (and thereby might also be candidates for dynamic signal encoding) can be found in [31].

In addition to the previously described but modest ultrasensitivity through oligomerisation [6, 7], we have seen that oligomerisation could also be an effective homeostatic regulatory mechanism to keep monomer concentration in a narrow range. In contrast to the dynamical phenomena, this more likely applies to proteins which are in a constitutive monomer/oligomer equilibrium. Recently, Frieden proposed oligomerisation as metabolic control mechanism [32]. Given that many enzymes oligomerise, monomer-homeostasis could be a good example. If enzyme function is inhibited e.g. by active site obstruction in an oligomeric complex [33], monomer-homeostasis could ensure a nearly constant performance of a metabolic activity over a wide range of total protein concentration (and therefore cellular conditions such as starvation or different cell cycle phases).

So far, few oligomeric proteins have been studied experimentally enough extensively to validate scenarios such as depicted in Figure 3B. Since individual species concentrations in the homeostatic scenario often differ by ≥ 2 orders of magnitude or more, experimental testing of such behaviour would at least require to determine the equilibrium distribution of monomeric and oligomeric species over several orders of magnitude of total protein concentration. Ideally, this would be complemented by kinetic data on oligomer (dis-)assembly. Both types of experiments can be technically challenging and likely need to be analysed through model fitting [34, 35].

### Combinatorial complexity

As the order of oligomers increases and/or PTMs are taken into account, the number of species and possible reactions quickly grows. This is a typical situation of ‘combinatorial explosion’ which poses a significant challenge for many signal transduction models [36, 37]. If PTMs are not considered and only one oligomeric species is relevant, oligomerisation pathways can be approximated via *generalised mass action* rate laws (i.e. power-laws) if the range total concentrations sufficiently restricted (data not shown).

Upon inclusion of PTMs, the combinatorial expansion of possible oligomerisation routes posed another unanticipated challenge: ensuring thermodynamic consistency of the model. Rate balancing offers a solution which is straightforward to apply to mass action kinetics models. An open question is how this procedure fares if PTMs do affect oligomerisation parameters. A plausible conjecture would be that once the balancing coefficients have been introduced into the rate equations, changing parameter values for individual reactions will not affect thermodynamic consistency.

For practical purposes, modelling higher-order oligomers with multiple PTM sites will generally require implicit modelling approaches. Rule based modelling, for instance, has been applied successfully for modelling EGF-receptor oligomerisation [38].

### Bistability

Ultrasensitivity and bistability are important properties in signal transduction networks for cellular decision making, allowing to respond in a switch-like, binary and sometimes irreversible fashion. Oligomerisation can also lead to ultrasensitivity and bistability by providing pseudo-multisite complexes (i.e. complexes with multiple identical PTM sites). Given previous work on ultrasensitivity and bistability arising through multi-site modification from the Kholodenko Lab and others [20, 22, 23, 25], this possibility seems obvious from a biochemical point of view, yet, has not been appreciated before. An interesting and unique twist of this motif is that the bistability resulting from modification of the oligomer extends to the monomer due to intrinsic substrate competition and because both species are in equilibrium with each other. We also demonstrated that kinetic cooperativity of multisite modification systems is not a requirement for bistability. If multiple enzymes regulate the modification steps and if some can only catalyse a subset of the individual modification steps, this leads effectively to the same kinetic asymmetry [23,25] as kinetic cooperativity. While oligomers might be particularly suited for this mechanism due their symmetrical structure, bistability through multi-enzyme regulation could in principle arise in any multisite PTM system.

The relevance of these findings is that they significantly expand the range of contexts in which one should look for biochemical ‘switches’ as both homo-oligomerisation and multi-enzyme regulation are extremely common. Phosphatases and GTPase activating proteins (GAPs), just to name two examples, are known to often act promiscuously on multiple substrates [39-41]. As a consequence many phosphorylation sites can be dephosphosphorylated by multiple phosphatases and many small GTPases (some of which can also dimerise), can be inactivated by multiple GAPs, creating potential situations in which bistability could occur. Alternatively, multi-enzyme regulation of the modification rather than demodification steps is also conceivable. Phosphorylating a protomer within a dimer, for example, could lead to a new binding site for a second kinase facilitating phosphorylation of the same residue in the other protomer. The combination of both mechanisms, oligomerisation and multi-enzyme regulation, therefore represents an interesting novel signalling motif that does not require feedback or kinetic cooperativity to generate bistable responses.

### Conclusion

We have demonstrated that homo-oligomers, making up approximately 30-50% of the proteome [1, 2], offer an even greater variety of regulatory mechanisms than previously appreciated. It is likely that some of these are relevant to many cellular signalling pathways. It also may partly explain why homo-oligomerisation is so commonly found throughout evolution. Hopefully, the presented findings will be helpful to modellers interested in homo-oligomeric signalling proteins and stimulate experimental research into signalling processes to which the presented findings might apply.

## Methods

Details on the computational procedures can be found in supplementary section 4.

## Acknowledgements

I thank Thomas Kampourakis, Stephen Martin, Franca Frater-nali and etra Gobbels for helpful discussions and critical reading of the manuscript and Brian Ingalls for his excellent textbook and general modelling advice. I acknowledge the British Heart Foundation for financial support (PhD studentship).

