## Supplementary Material for "Homo-Oligomerisation in Signal Transduction: Dynamics, Homeostasis, Ultrasensitivity, Bistability"

Daniel Koch
orcid.org/0000-0002-4893-1774

November 2019

---

#### Contents

**1 Reaction rates and ODEs for trimerisation model including PTMs** **2**

**2 Balancing the rates of the trimerisation model including PTMs** **3**

**3 Proof: conservation of oligomerisation rates** **5**

**4 Computational methods** **8**

**5 Supplementary figures** **8**

**6 References** **9**

---

### 1 Reaction rates and ODEs for trimerisation model including PTMs

This section contains the reaction rates and ODEs for the reaction scheme presented in Figure 4B of the main text.

##### Reaction rates

association & dissociation reactions

$$v7 = k_7[A][AA] \qquad v8 = k_8[AAA]$$
$$v9 = k_9[A^*][AA] \qquad v10 = k_{10}[AAA^*]$$
$$v11 = k_{11}[A][AA^*] \qquad v12 = k_{12}[AAA^*]$$
$$v13 = k_{13}[A^*][AA^*] \qquad v14 = k_{14}[AAA^{**}]$$
$$v15 = k_{15}[A][AA^{**}] \qquad v16 = k_{16}[AAA^{**}]$$
$$v17 = k_{17}[A^*][AA^{**}] \qquad v18 = k_{18}[AAA^{***}]$$
$$v23 = k_{23}[A]^2 \qquad v24 = k_{24}[AA]$$
$$v25 = k_{25}[A][A^*] \qquad v26 = k_{26}[AA^*]$$
$$v27 = k_{27}[A^*]^2 \qquad v28 = k_{28}[AA^{**}]$$

modification reactions

$$v1 = \frac{k_1[E1][AAA]}{K_1(1 + \frac{[A]}{K_{29}} + \frac{[AA]}{K_{19}} + \frac{[AA^*]}{K_{21}} + \frac{[AAA^*]}{K_3} + \frac{[AAA^{**}]}{K_5}) + [AAA]}$$
$$v3 = \frac{k_3[E1][AAA^*]}{K_3(1 + \frac{[A]}{K_{29}} + \frac{[AA]}{K_{19}} + \frac{[AA^*]}{K_{21}} + \frac{[AAA]}{K_1} + \frac{[AAA^{**}]}{K_5}) + [AAA^*]}$$
$$v5 = \frac{k_5[E1][AAA^{**}]}{K_5(1 + \frac{[A]}{K_{29}} + \frac{[AA]}{K_{19}} + \frac{[AA^*]}{K_{21}} + \frac{[AAA]}{K_1} + \frac{[AAA^*]}{K_3}) + [AAA^{**}]}$$
$$v19 = \frac{k_{19}[E1][AA]}{K_{19}(1 + \frac{[A]}{K_{29}} + \frac{[AA^*]}{K_{21}} + \frac{[AAA]}{K_1} + \frac{[AAA^*]}{K_3} + \frac{[AAA^{**}]}{K_5}) + [AA]}$$
$$v21 = \frac{k_{21}[E1][AA^*]}{K_{21}(1 + \frac{[A]}{K_{29}} + \frac{[AA]}{K_{19}} + \frac{[AAA]}{K_1} + \frac{[AAA^*]}{K_3} + \frac{[AAA^{**}]}{K_5}) + [AA^*]}$$
$$v29 = \frac{k_{29}[E1][A]}{K_{29}(1 + \frac{[AA]}{K_{19}} + \frac{[AA^*]}{K_{21}} + \frac{[AAA]}{K_1} + \frac{[AAA^*]}{K_3} + \frac{[AAA^{**}]}{K_5}) + [A]}$$

demodification reactions

$$v2 = \frac{k_2[E2][AAA^*]}{K_2(1 + \frac{[A^*]}{K_{30}} + \frac{[AA^*]}{K_{20}} + \frac{[AA^{**}]}{K_{22}} + \frac{[AAA^{**}]}{K_4} + \frac{[AAA^{***}]}{K_6}) + [AAA^*]}$$
$$v4 = \frac{k_4[E2][AAA^{**}]}{K_4(1 + \frac{[A^*]}{K_{30}} + \frac{[AA^*]}{K_{20}} + \frac{[AA^{**}]}{K_{22}} + \frac{[AAA^*]}{K_2} + \frac{[AAA^{***}]}{K_6}) + [AAA^{**}]}$$
$$v6 = \frac{k_6[E2][AAA^{***}]}{K_6(1 + \frac{[A^*]}{K_{30}} + \frac{[AA^*]}{K_{20}} + \frac{[AA^{**}]}{K_{22}} + \frac{[AAA^*]}{K_2} + \frac{[AAA^{**}]}{K_4}) + [AAA^{***}]}$$
$$v20 = \frac{k_{20}[E2][AA^*]}{K_{20}(1 + \frac{[A^*]}{K_{30}} + \frac{[AA^{**}]}{K_{22}} + \frac{[AAA^*]}{K_2} + \frac{[AAA^{**}]}{K_4} + \frac{[AAA^{***}]}{K_6}) + [AA^*]}$$
$$v22 = \frac{k_{22}[E2][AA^{**}]}{K_{22}(1 + \frac{[A^*]}{K_{30}} + \frac{[AA^*]}{K_{20}} + \frac{[AAA^*]}{K_2} + \frac{[AAA^{**}]}{K_4} + \frac{[AAA^{***}]}{K_6}) + [AA^{**}]}$$
$$v30 = \frac{k_{30}[E2][A^*]}{K_{30}(1 + \frac{[AA^*]}{K_{20}} + \frac{[AA^{**}]}{K_{22}} + \frac{[AAA^*]}{K_2} + \frac{[AAA^{**}]}{K_4} + \frac{[AAA^{***}]}{K_6}) + [A^*]}$$

The ODE system is:

$$\frac{d}{dt}[A] = v8 + v12 + v16 + 2 \cdot v24 + v26 + v30 - v7 - v11 - v15 - 2 \cdot v23 - v25 - v29$$
$$\frac{d}{dt}[A^*] = v10 + v14 + v18 + 2 \cdot v28 + v26 + v29 - v9 - v13 - v17 - 2 \cdot v27 - v25 - v30$$
$$\frac{d}{dt}[AA] = v8 + v10 + v20 + v23 - v7 - v9 - v19 - v24$$
$$\frac{d}{dt}[AA^*] = v12 + v14 + v19 + v22 + v25 - v11 - v13 - v20 - v21 - v26$$
$$\frac{d}{dt}[AA^{**}] = v16 + v18 + v21 + v27 - v15 - v17 - v22 - v28$$
$$\frac{d}{dt}[AAA] = v2 + v7 - v1 - v8$$
$$\frac{d}{dt}[AAA^*] = v1 + v4 + v9 + v11 - v2 - v3 - v10 - v12$$
$$\frac{d}{dt}[AAA^{**}] = v3 + v6 + v13 + v15 - v4 - v5 - v14 - v16$$
$$\frac{d}{dt}[AAA^{***}] = v5 + v17 - v6 - v18$$

#### 2 Balancing the rates of the trimerisation model including PTMs

As for the dimer model, we work under the assumption that all association steps leading to an oligomeric complex $A_n$ have the same rate constant $k_i$ and all dissociation steps have the same rate constant $k'_i$. As trimers can only be formed by association of dimers and monomers (i.e. $I(A_3) = \{\{1,2\}\}$), we leave out the index $i \in I$ for convenience. We begin the rate balancing of the trimerisation model by noticing that the lower part of the trimerisation scheme depicted in Figure 4B is structurally identical to the dimerisation scheme in Figure 4A. We thus can use the same rate balancing coefficients for the dimerisation steps in the trimer model, i.e. we balance $v25$ by multiplying it with 2 to obtain $v25 = 2k[A][A^*]$. We next apply the rate conservation principle to the formation of trimers and obtain:

$$v7 + v9 + v11 + v13 + v15 + v17 = k[A_t][AA_t] = k([A] + [A^*]) \cdot ([AA] + [AA^*] + [AA^{**}])$$

→ *expanding RHS, substituting LHS:*

$$k[AA][A] + k[AA][A^*] + k[AA^*][A] + k[AA^*][A^*] + k[AA^{**}][A] + k[AA^{**}][A^*]$$
$$= k[AA][A] + k[AA][A^*] + k[AA^*][A] + k[AA^*][A^*] + k[AA^{**}][A] + k[AA^{**}][A^*]$$

→ *no balancing necessary*

→ *assigning reaction rates:*

$$v7 = k[AA][A]$$
$$v9 = k[AA][A^*]$$
$$v11 = k[AA^*][A]$$
$$v13 = k[AA^*][A^*]$$
$$v15 = k[AA^{**}][A]$$
$$v17 = k[AA^{**}][A^*]$$

Applying the rate conservation principle to the dissociation of trimers we obtain:

$$v8 + v10 + v12 + v14 + v16 + v18 = k'[AAA_t] = k'([AAA] + [AAA^*] + [AAA^{**}] + [AAA^{***}])$$

→ *expanding RHS, substituting LHS:*

$$k'[AAA] + k'[AAA^*] + k'[AAA^*] + k'[AAA^{**}] + k'[AAA^{**}] + k'[AAA^{***}]$$
$$= k'[AAA] + k'[AAA^*] + k'[AAA^{**}] + k'[AAA^{***}]$$

We notice that the dissociation rates for singly and dually modified trimers appear twice in the LHS, but only once in the RHS. We therefore need to
→ *balance deviating terms in LHS:*

$$k'[AAA] + \mathbf{\frac{1}{2}}k'[AAA^*] + \mathbf{\frac{1}{2}}k'[AAA^*] + \mathbf{\frac{1}{2}}k'[AAA^{**}] + \mathbf{\frac{1}{2}}k'[AAA^{**}] + k'[AAA^{***}]$$
$$= k'[AAA] + k'[AAA^*] + k'[AAA^{**}] + k'[AAA^{***}]$$

→ *assigning reaction rates:*

$$v8 = k'[AAA]$$
$$v10 = \frac{1}{2}k'[AAA^*]$$
$$v12 = \frac{1}{2}k'[AAA^*]$$
$$v14 = \frac{1}{2}k'[AAA^{**}]$$
$$v16 = \frac{1}{2}k'[AAA^{**}]$$
$$v18 = k'[AAA^{***}]$$

### 3 Proof: conservation of oligomerisation rates

To show the validity of the principle of conservation of oligomerisation rates, we require the principle of detailed balance. While it has often been treated as a fundamental postulate, it can be derived from microscopic reversibility in physics, which is why we treat it as a lemma. [2,3]

**Lemma: principle of detailed balance**

> "When a system is at equilibrium this means that the forward rate of a molecular process has to be equal to the reverse rate of that process. Applying this to a chemical reaction observed on a macroscopic scale means that at equilibrium, the forward rate of each step is equal to the reverse rate of that step; this is the principle of detailed balance." [4]

For a system in which some molecular species $S$ can be formed by $j \in \mathbb{N}^{\geq 1}$ reversible reactions this means at equilibrium:

$$\sum_{i=1}^{j} v_i = \sum_{i=1}^{j} v_i', \tag{1}$$

where $v_i$ is the forward rate and $v_i'$ the reverse rate of reaction $i$.

Although the multitude sums might give the impression of being complicated, the idea for the principle's proof itself is simple: applying the principle of detailed balance and the assumption that the PTM does not influence oligomerisation (expectation (e) from the main text) to the n-tamer equilibrium concentrations results in the same equations as given by the postulated conservation principle but restricted to equilibrium. We can then prove the principle by contradiction. If we assume that the conservation principle would be false and apply the resulting inequalities to the equilibrium situation, we obtain direct contradictions to the before derived equations describing the equilibrium. For convenience, the principle to be proven is repeated in summarized form below:

**Conservation of oligomerisation rates**

Let $A_n$ be an oligomeric complex with $n \in \mathbb{N}^{\geq 1}$ protomers[1] which can be formed through $j \in \mathbb{N}^{\geq 1}$ reversible bimolecular association reactions between lower-order complexes $A^p_{r_i}$ and $A^q_{s_i}$, where $p + q \leq n$ and $r_i + s_i = n$, $i \in I(A_n)$. The PTM does not influence the oligomerisation reaction.

It follows that at all times, the sum of all rates $v_i$, $1 \leq i \leq j$, of reactions leading to $A_n$ of any modification status is the association rate based on the total concentrations (i.e. all modification isoforms) of $A_n$'s educts. The sum of all rates $v'_i$, $1 \leq i \leq j$, of reactions dissociating $A_n$ of any modification status is equal to the dissociation rate based on $A_n$'s total concentration. That is

$$\sum_{i=1}^{j} v_i = \sum_{i \in I} (k_i \cdot \sum_{m=0}^{r_i} [A^m_{r_i}] \cdot \sum_{m=0}^{s_i} [A^m_{s_i}]), \tag{2}$$

and

$$\sum_{i=1}^{j} v'_i = \sum_{i \in I} (k'_i \cdot \sum_{m=0}^{n} [A^m_n]). \tag{3}$$

**Proof:** For the construction of the contradiction it is necessary to consider the situation at equilibrium. By assumption, the PTM does not influence oligomerisation. From (a1) and (a2) it follows that all forward steps between $A^p_{r_i}$ and $A^q_{s_i}$ producing $A^m_n$ have the same association rate constant $k_i$ and each corresponding reverse step has the same dissociation rate constant $k'_i$ for all possible values of $m, p$ and $q$. According to (d), equilibrium is only determined by the total concentration of each oligomeric complex, i.e. the relevant educt concentrations are $\sum_{m=0}^{r_i} [A^m_{r_i}]_{eq}$ and $\sum_{m=0}^{s_i} [A^m_{s_i}]_{eq}$, whereas the relevant product concentration is $\sum_{m=0}^{n} [A^m_n]_{eq}$. Thus, the expressions for forward and reverse rates are

$$k_i \cdot \sum_{m=0}^{r_i} [A^m_{r_i}]_{eq} \cdot \sum_{m=0}^{s_i} [A^m_{s_i}]_{eq}$$

and

$$k'_i \cdot \sum_{m=0}^{n} [A^m_n]_{eq},$$

respectively, for all $i \in I$. Applying the principle of detailed balance (1) to the formation and dissociation of $A_n$ at equilibrium and substituting above rate expressions therefore gives:

$$\sum_{i=1}^{j} v_i = \sum_{i \in I} (k_i \cdot \sum_{m=0}^{r} [A^m_{r_i}]_{eq} \cdot \sum_{m=0}^{s} [A^m_{s_i}]_{eq}) = \sum_{i \in I} (k'_i \cdot \sum_{m=0}^{n} [A^m_n]_{eq}) = \sum_{i=1}^{j} v'_i. \tag{4}$$

[1] Monomers are considered as oligomers for purely formal reasons.

Separating (4) into its first and last identity yields

$$\sum_{i=1}^{j} v_i = \sum_{i \in I} (k_i \cdot \sum_{m=0}^{r} [A_{r_i}^m]_{eq} \cdot \sum_{m=0}^{s} [A_{s_i}^m]_{eq}), \tag{5}$$

and

$$\sum_{i=1}^{j} v_i' = \sum_{i \in I} (k_i' \cdot \sum_{m=0}^{n} [A_n^m]_{eq}). \tag{6}$$

Let us for a moment assume that the conservation of oligomerisation rates would not be true. We have to distinguish three cases: equation (2) is false, equation (3) is false, both (2) and (3) are false.

**Case 1:** equation (2) is false, i.e.

$$\sum_{i=1}^{j} v_i \neq \sum_{i \in I} (k_i \cdot \sum_{m=0}^{r} [A_{r_i}^m] \cdot \sum_{m=0}^{s} [A_{s_i}^m]),$$

$$\text{at equilibrium} \quad \sum_{i=1}^{j} v_i \neq \sum_{i \in I} (k_i \cdot \sum_{m=0}^{r} [A_{r_i}^m]_{eq} \cdot \sum_{m=0}^{s} [A_{s_i}^m]_{eq}),$$

in contradiction to (5).

**Case 2:** equation (3) is false, i.e.

$$\sum_{i=1}^{j} v_i' \neq \sum_{i \in I} (k_i' \cdot \sum_{m=0}^{n} [A_n^m]),$$

$$\text{at equilibrium} \quad \sum_{i=1}^{j} v_i' \neq \sum_{i \in I} (k_i' \cdot \sum_{m=0}^{n} [A_n^m]_{eq}),$$

in contradiction to (6).

**Case 3:** Both (2) and (3) are false. The contradiction follows from case 1 and case 2.

□

### 4 Computational methods

Presented models have been implemented as MATLAB® scripts for numerical simulation and model analysis. All simulations have been performed using the ode23s integrator. Sensitivity analysis has been performed as described in [1]. The model code will be released on a public database upon final publication.

Bifurcation diagrams have been generated using a custom algorithm that iteratively identifies the unstable steady states. If the algorithm does not converge within the specified number of iterations, it approximates the unstable steady state. The approximation is based on the distribution of concentrations from all time courses taking advantage of the fact that simulations very close to the unstable steady state take a long time until they eventually tip to either of the stable steady states. For more details, please refer to: https://www.ebi.ac.uk/biomodels/MODEL1910220002.

### 5 Supplementary figures

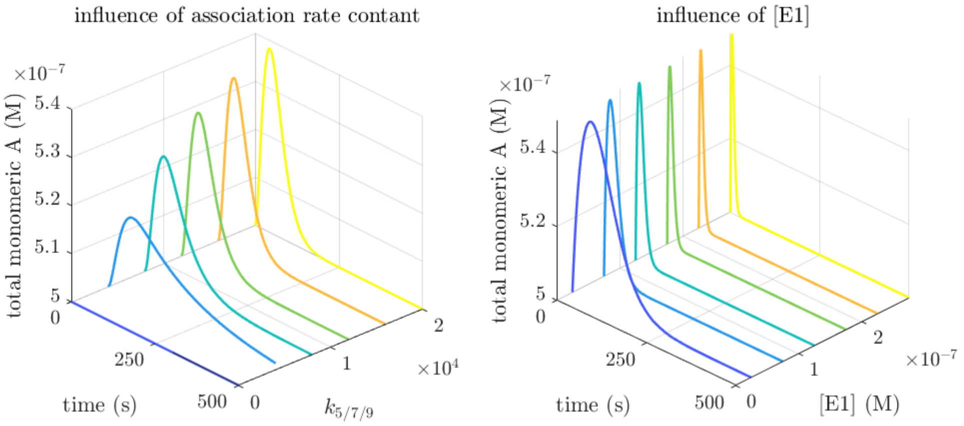


**Figure S1**
Dependence of transient changes in the mass action dimerisation model on association rate constants and concentration of the modifying enzyme.

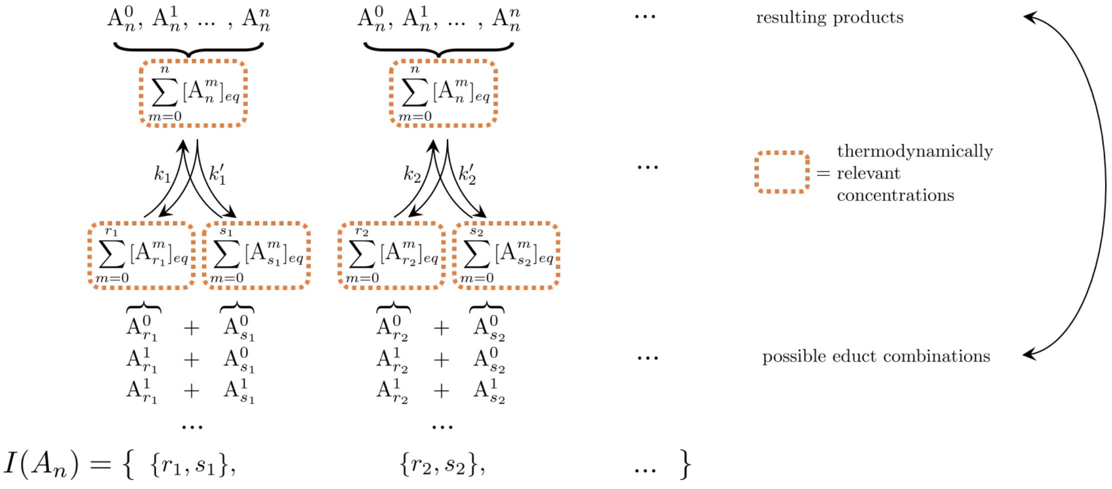


**Figure S2**
Illustration of the n-tamer equilibrium situation if a PTM does not influence the oligomerisation reaction. According to expectation (d) in the main text, we expect the equilibrium to be solely determined by the total n-tamer concentrations $\sum_{m=0}^{n}[A_n^m]_{eq}$ if the PTM status does not influence oligomerisation. The set $I(A_n)$ denotes the actually occurring combinations of oligomeric orders of $A_n$'s educts, i.e. $|I(A_n)|$ is the number of reversible reactions that would produce/consume $A_n$ if PTMs were not taken into consideration.
